# The TALE effector PthA4 of *Xanthomonas citri* subsp. *citri* indirectly activates an expansin gene *CsEXP2* and an endoglucanase *CsEG1* via CsLOB1 to cause citrus canker symptoms

**DOI:** 10.1101/2025.03.20.644280

**Authors:** Rikky Rai, Nian Wang

**Author notes:** Corresponding author: Nian Wang.

## Abstract

Citrus canker caused by *Xanthomonas citri* subsp. citri (Xcc) is an important citrus disease worldwide. *PthA4* is the most important pathogenicity gene of Xcc and encodes a transcription activator like effector (TALE) secreted by the type III secretion system. PthA4 is known to activate the expression of *CsLOB1*, the canker susceptibility gene and a transcription factor, to cause citrus canker symptoms. Extensive effort was made to identify downstream targets of CsLOB1 to investigate the mechanism underlying canker symptom development. However, none of identified CsLOB1 target genes have been confirmed to be involved in citrus canker development. Here, we first identified the direct targets of CsLOB1 by generating promoter-*uidA* (GUS) reporter fusion construct for the 13 genes highly induced by both PthA4 and CsLOB1 and monitored the reporter activity in *N. benthamiana* leaves co-expressing *CsLOB1*. *Agrobacterium tumefaciens*-mediated transient expression of *CsLOB1* activated seven gene promoters in *N. benthamiana* including Cs7g18460, Orange1.1t00600, Cs6g17190, Cs7g32410 (*CsEXP2*), Cs2g27100, Cs2g20750 (*CsEG1*), and Cs9g17380. Next, we constructed dTALEs to target unique sequences in the promoters of the seven direct target genes of CsLOB1 and transformed them into Xcc*pthA4*::Tn5 mutant. Our results indicate that a combination of 5 and 7 dTALEs caused canker-like symptoms in the inoculated citrus leaves. In addition, dTALECsEXP2 and dTALECsEG1 caused water soaking and pustules, which are typical canker symptoms. Taken together, Xcc indirectly activates CsEXP2 and CsEG1 via PthA4-CsLOB1 to cause canker symptoms.

**Author summary:** Ptha4 is responsible for the canker symptoms of Xcc and activate the expression of *CsLOB1* of citrus to cause the canker symptoms. Extensive effort by multiple groups has been made to identify downstream targets of CsLOB1 to understand the mechanism underlying canker symptom development. However, none of the CsLOB1 targets have been shown to cause canker symptoms. Here we have demonstrated that Xcc indirectly activates CsEXP2, an expansin gene, and CsEG1, an endoglucanase, via PthA4-CsLOB1 to cause canker symptoms. Identification of direct targets of CsLOB1 provides alternative target genes for genetic improvement of citrus against canker via genome editing.

## Introduction

Citrus canker caused by *Xanthomonas citri* subsp. *citri* (Xcc) is one of the most severe citrus diseases worldwide. It is endemic in most citrus production regions including Brazil, China, India, Mexico, USA except Australia and the Mediterranean region [1]. PthA4, the most important pathogenicity factor of Xcc, is a transcription activator like effector (TALE) which is secreted by the type III secretion system. PthA4 is responsible for the canker symptom development including water soaking, hypertrophy and hyperplasia and further release and spread from the lesions [2, 3]. PthA4 binds to the promoter region (effector binding element (EBE)) to activate the expression of *CsLOB1*, the canker susceptibility gene, thus causing canker symptoms [4]. Natural variations in the EBE region of *LOB1* gene in *Atalantia buxifolia* contributes to its resistance against Xcc [5]. Similarly, TALEs have been known to activate disease susceptibility genes including *OsSULTR3*:6 [6], *UPA20* [7], and *SWEET* [8, 9] to promote pathogen growth or symptom development in many pathosystems.

Diverse approaches have been developed to control plant diseases by targeting the susceptibility genes. Genome editing of the EBE region of susceptibility genes has been widely used for engineering disease resistant crops. For instance, genome editing of the EBE of *SWEET* genes has led to the development of disease resistant rice varieties against bacterial blight [10, 11]. Genome editing of the EBE region or TATA box of *CsLOB1* genes has increased disease resistance against citrus canker [12–19]. Gene silencing of *CsLOB1* via both transgenic approach [20] and application of antisense oligonucleotide [21] increase disease resistance against citrus canker. Importantly, non-transgenic genome editing of the EBE of *CsLOB1* has led to the development of canker resistant citrus varieties that have been approved for commercialization by two federal regulatory agencies The Animal and Plant Health Inspection Service (APHIS) and Environmental Protection Agency (EPA) in the USA [18, 22–24].

CsLOB1 belongs to the lateral organ boundaries domain (LBD) family transcription factors [25, 26]. LBD proteins have diverse functions in plant growth and development [27] including in lateral root formation [28], and microspore division [29]. Extensive effort has been made to identify downstream targets of CsLOB1 to understand the mechanism underlying canker symptom development. Duan et al. conducted bioinformatics and electrophoretic mobility shift assays and showed that CsLOB1 binds to the promoter of Cs2g20600, a zinc finger C3HC4-type RING finger gene [30]. Zou et al. identified 565 putative CsLOB1-targeted genes via chromatin immunoprecipitation-sequencing (Chip-seq) [20]. de Souza-Neto et al. reported that CsLOB1, but not PthA4, directly actives the expression of an expansin gene *CsLIEXP1* (orange1.1t00187) [31]. Long et al. showed that CsLOB1 promoted the expression of *CsNCED1*-1, which encodes 9-cis-epoxycarotenoid dioxygenase, an enzyme in abscisic acid biosynthesis, by binding to its promoter. Chen et al. reported that CsLOB1 binds to the promoter of Cs9g12620, which encode a carbohydrate-binding protein [32]. In addition, it was reported that CsLOB1 activates a cellulase gene *CsCEL20* by targeting its promoter region [33]. However, none of the previous studies have established that the identified CsLOB1 target genes are involved in citrus canker development. In this study, we cracked this puzzle using designer TALEs to activate putative CsLOB1 target genes. We have shown that CsLOB1 directly activates seven genes and activation of Cs7g32410 (hereafter *CsEXP2*), which encodes an expansin protein, and Cs2g20750 (hereafter *CsEG1*), which encodes an endoglucanase protein, using designer TALEs, led to canker symptoms.

## Results

### Identification of candidate direct target genes of CsLOB1

To identify the candidate target genes of CsLOB1 that are involved in causing citrus canker, we reasoned that genes whose expression was dependent on the presence of PthA4 from *Xcc* or CsLOB1 of citrus are likely to be regulated by CsLOB1. To identify such genes, we analyzed gene expression datasets and selected 14 candidate genes that are commonly and highly upregulated by both PthA4 and CsLOB1 [4, 20, 25, 34] including Cs1g16560, Cs7g32410, Cs6g17190, Cs2g27100, Cs9g17380, Cs2g27090, Cs3g18550, Cs2g21440, Cs7g18460, Cs5g33680, orange1.1t00600, Cs2g05560, Cs2g20750, and Cs2g20600 (Supplementary Table 1) [4, 20, 25, 34].

### Seven candidate genes displayed CsLOB1-dependent promoter activity

To examine if these 14 genes were directly regulated by CsLOB1, we amplified their promoter regions (approximately 1 kb in length) and checked their promoter activity in the presence or absence of CsLOB1 as described previously [31]. We were unable to amplify the promoter region of Cs3g18550 despite multiple attempts, so we proceeded with the promoter regions of the 13 other genes. We generated promoter-*uidA* (ß-glucuronidase (GUS) reporter fusion construct for the 13 promoters and monitored the reporter activity in *N. benthamiana* leaves transiently expressing CsLOB1 or EV (control) (Fig.1) [31]. The 13 promoter::GUS fusions (pCs1g16560::*uidA*, pCs7g32410::*uidA*, pCs6g17190::*uidA*, pCs2g27100::*uidA*, pCs9g17380::*uidA*, pCs2g27090::*uidA*, pCs2g21440::*uidA*, pCs7g18460::*uidA*, pCs5g33680::*uidA*, orange1.1t00600::*uidA*, pCs2g05560::*uidA*, pCs2g20750::*uidA*, and pCs2g20600::*uidA*) were individually co-expressed in *N. benthamiana* with the 35S::CsLOB1-HA plasmid by *Agrobacterium*-mediated transformation (Fig.1 B). As a positive control, the empty vector with 35S promoter upstream of *uidA* in pCAMBIA1380 was co-expressed in *N. benthamiana* in the presence or absence of CsLOB1 (Fig.1C). The co-inoculation of the 35S promoter fused to GUS resulted in no significant change (P ≤ 0.05) in GUS expression in the presence or absence of the CsLOB1 (Fig. 1D). Our results showed that *A. tumefaciens*-mediated transient ectopic expression of *CsLOB1* activated seven gene promoters in *N. benthamiana* at 48 hours post inoculation (hpi) (Fig.1D) including Cs7g18460, Orange1.1t00600, Cs6g17190, Cs7g32410, Cs2g27100, Cs2g20750, and Cs9g17380, but not the six other genes. Cs7g18460, Orange1.1t00600, Cs6g17190, Cs7g32410, Cs2g27100, Cs2g20750, and Cs9g17380 encode glucomannan 4-beta-mannosyltransferase 2, 3-oxo-5-alpha-steroid 4-dehydrogenase, gibberellin-regulated protein 5, expansin, GDSL esterase/lipase, endoglucanase 11, and uncharacterized protein, respectively (Supplementary Table 1). These results indicate that CsLOB1 directly induces the expression of these seven genes in *N. benthamiana* by potentially targeting an EBE site in their promoter regions. The seven promoters were searched for the 6-bp LBD motif (GCGGCG), which is known to bind to LOB domain proteins [35, 36]. Among the 7 induced promoters by CsLOB1, Cs6g17190, Cs9g17380, Orange1.1t00600 but not Cs7g18460, Cs7g32410, Cs2g27100 and Cs2g20750 contain the LBD motif, suggesting other motifs can interact with LOB domain proteins.

**Fig. 1.**
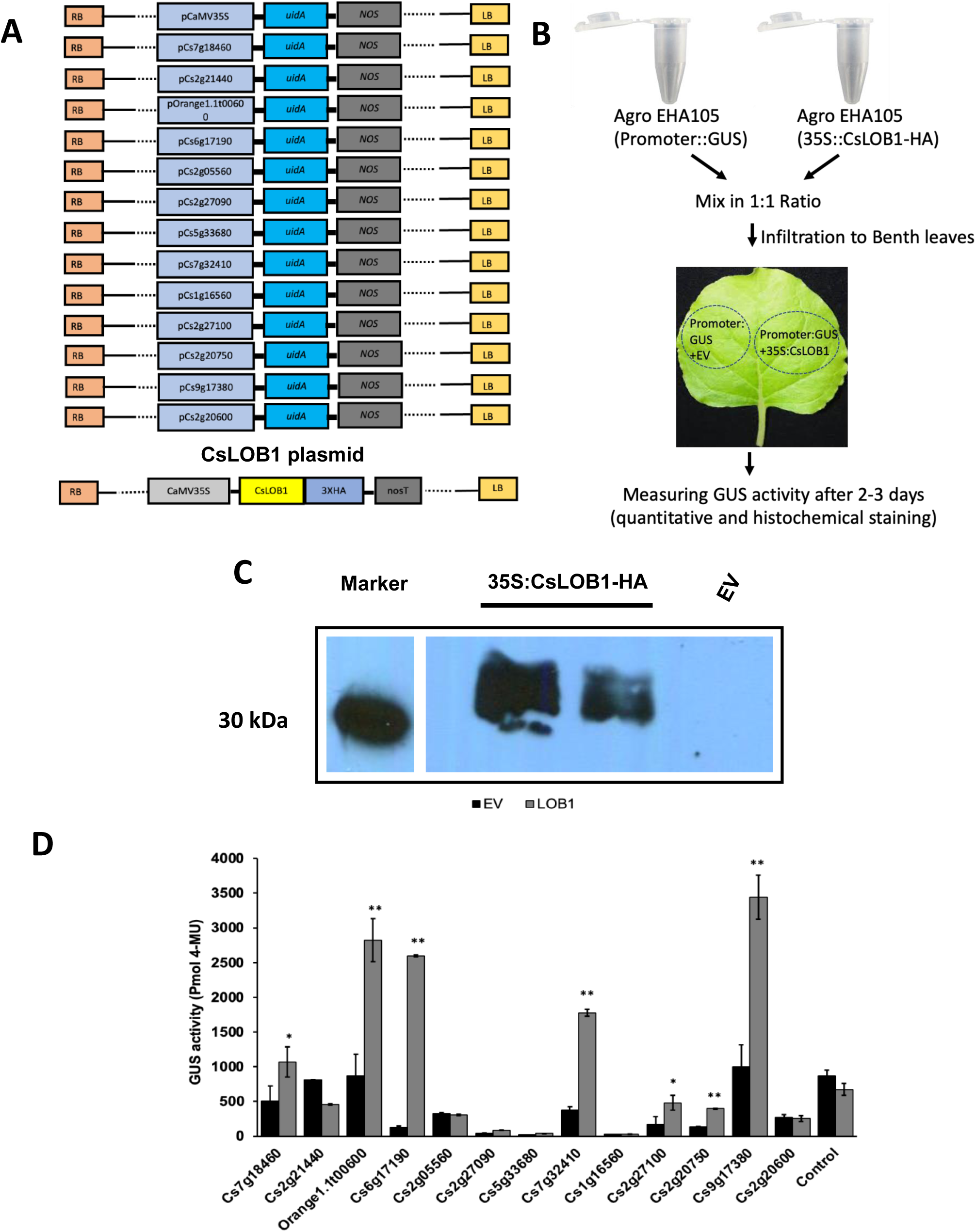
Identification of genes directly regulated by CsLOB1. A. Schematic diagram of reporter and CsLOB1 expression constructs. Reporter constructs contained the promoter of genes of interest fused to the GUS reporter. CsLOB1 expression construct is driven by the 35S promoter and fused to HA tag. B. Schematic representation of *Agrobacterium* mediated inoculation of reporter and expression construct in *Nicotiana benthamiana* (*Nb*) leaves to measure the promoter activity in the presence or absence of CsLOB1. C. Detection of CsLOB1 expression in *Nb* leaves by western blotting. D. GUS assay. Samples were collected 48 hpi and GUS activity was measured three times with similar results. Promoters of the seven genes (Cs7g18460, Orange1.1t00600, Cs6g17190, Cs7g32410, Cs2g27100,Cs2g20750, and Cs9g17380) showed induction when CsLOB1 was ectopically expressed compared to empty vector (EV). The 35S promoter fused with GUS was used as a control. Error bars indicate means ± SD (*n* = 3), and asterisks indicate significant differences (∗*P* ≤ 0.05; ∗∗*P* ≤ 0.01).

### Artificial dTALEs activate the expression of the identified CsLOB1 targets and trigger canker symptoms

Next, we tested whether activation of Cs7g18460, Orange1.1t00600, Cs6g17190, Cs7g32410, Cs2g27100, Cs2g20750, or Cs9g17380 can cause citrus canker symptoms. For this purpose, designer TALEs (dTALEs) have been used to activate genes of interest specifically [37, 38]. Accordingly, we constructed dTALEs to target unique sequences in the promoters of the seven direct target genes of CsLOB1 to induce each gene individually using optimized repeat variable di-amino acid (RVD) residues (Fig. 2). The dTALEs were cloned into a pBBR5 vector with a C-terminal HA-tag epitope and introduced into Xcc306*pthA4*::Tn5 and tested whether they can cause citrus canker symptoms on citrus leaves. Western blot analysis confirmed that the dTALEs were expressed in Xcc (Fig.2C). RT-qPCR assays showed that Cs2g27100, Cs7g18460, Orange1.1t00600, Cs6g17190, Cs7g32410, and Cs2g20750 genes were significantly induced by their respective dTALEs whereas Cs9g17380 induction was not statistically significant (Fig.3). In all cases, the 7 genes were highly induced in the positive controls *Xcc* WT and *pthA4*::Tn5 mutant transformed with dTALECsLOB1 (Fig. 4). We also checked whether dTALEs targeting genes of interest impact the expression of CsLOB1 by RT-qPCR. Expectedly, *CsLOB1* was activated by *Xcc* WT and dTALECsLOB1 compared to the *pthA4* mutant. None of the dTALEs targeting the promoters of the 7 genes induced the expression of *CsLOB1*, eliminating the potential interference from CsLOB1 (Fig. 4).

**Fig. 2.**
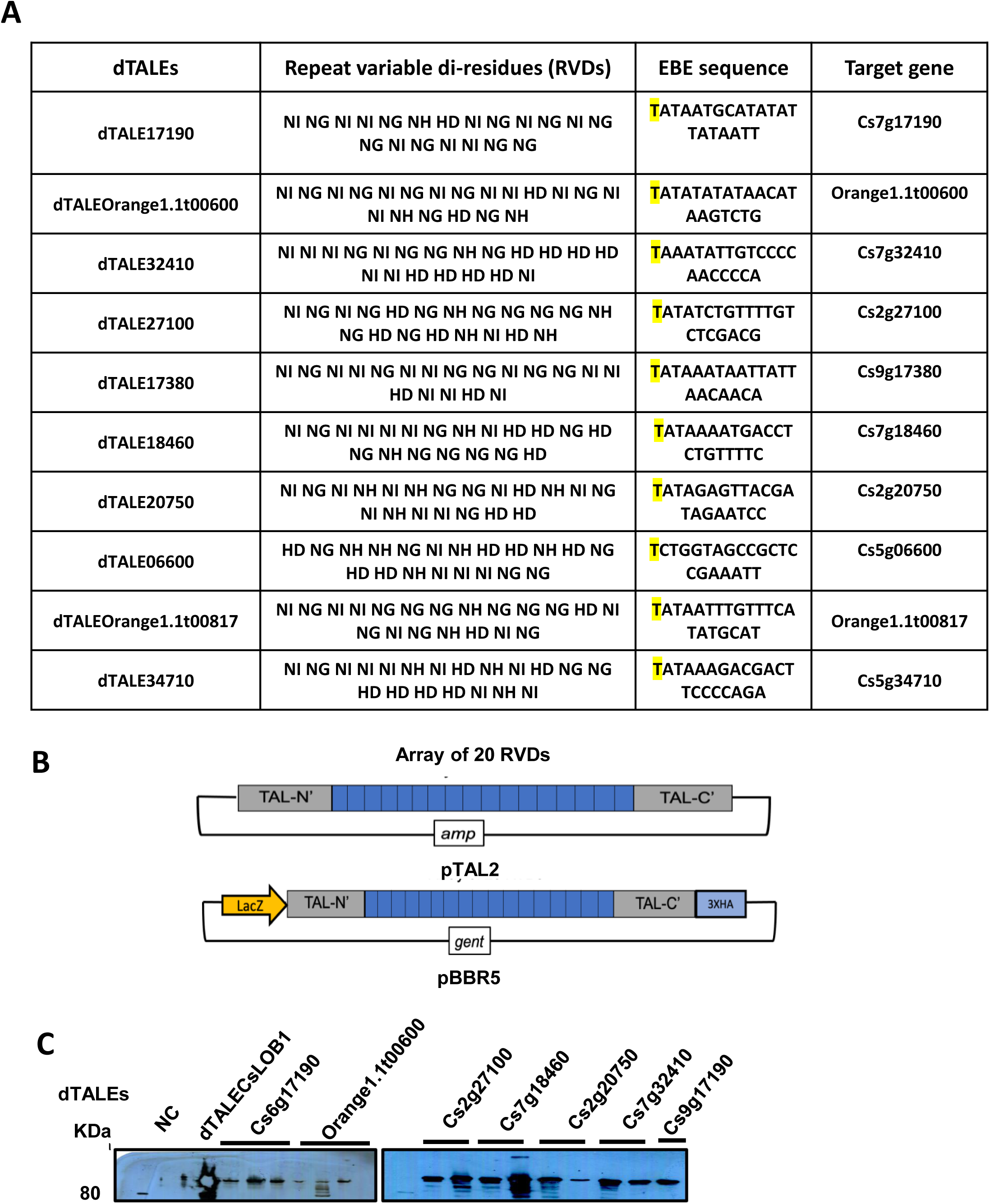
Activation of putative CsLOB1 target genes using designer TALEs (dTALEs). A. RVD repeat arrays of dTALEs used to activate the target genes. B. dTALE constructs were digested from pTAL2 destination vector, subcloned into pBBR5 with an HA tag at C-terminal. C. Detection of expression of dTALEs using Western Blotting. Total protein was extracted from overnight cultures of *Xcc pthA4*:Tn5 [Negative control (NC)], and *Xcc pthA4*:Tn5 transformed with the dTALE targeting CsLOB1 as positive control and dTALEs targeting other genes. Samples were separated by SDS PAGE and immunoblotted with the anti-HA antibody. Multiple lanes under the black line indicate independent colonies transformed with the respective dTALEs constructs.

**Fig. 3.**
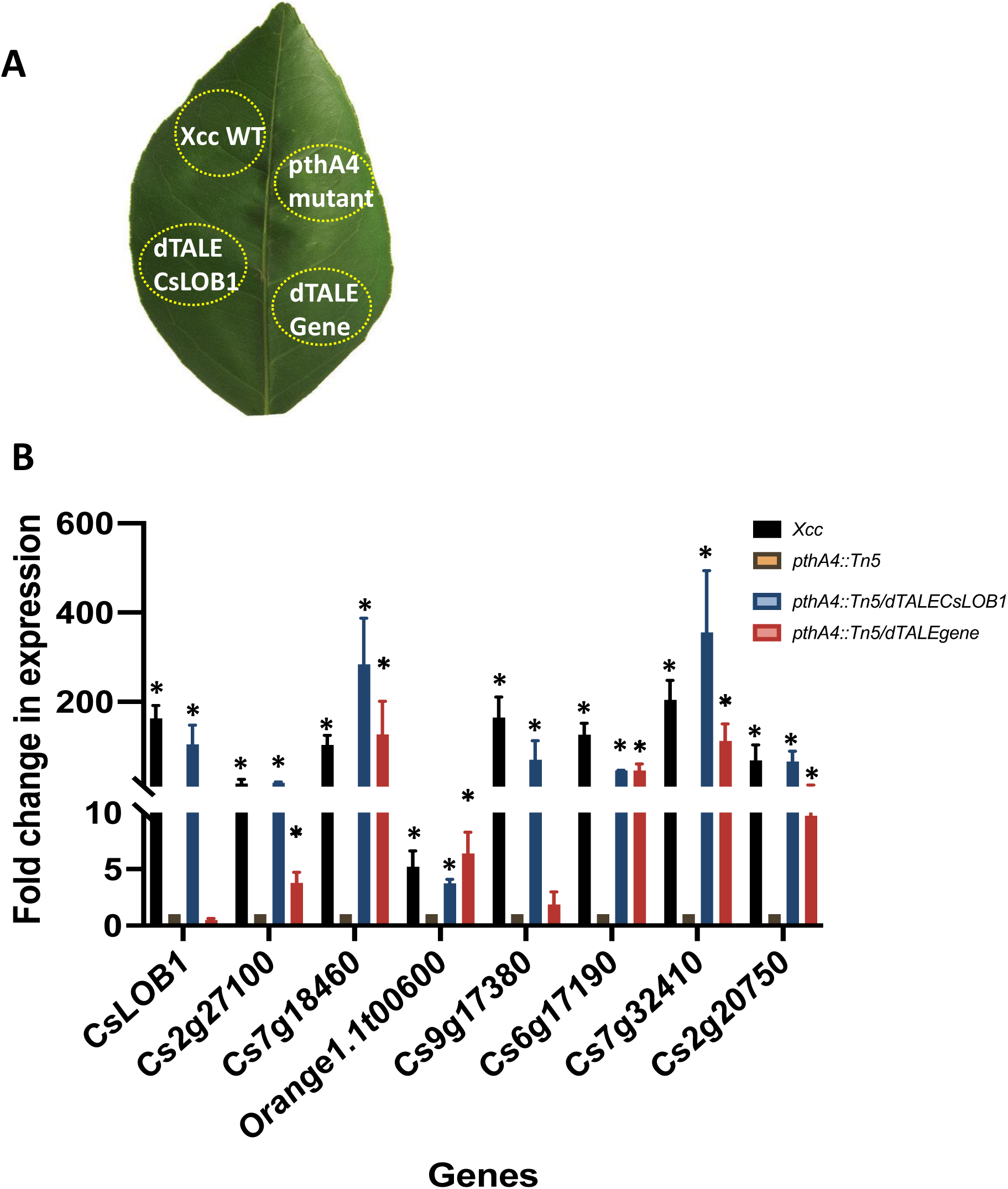
Confirmation of the expression of putative CsLOB1 target genes driven by dTALEs. A, A schematic representation of Valencia sweet orange leaf infiltrated with *Xcc* strains WT, Xcc*pthA4*::Tn5 mutant, and Xcc*pthA4*::Tn5 strains carrying dTALE*CsLOB1* or dTALE gene of interest using a needleless syringe. B. Expression analysis of *CsLOB1*, *Cs2g27100*, *Cs7g18460*, *Orange1.1t00600*, *Cs9g17380*, *Cs6g17190*, *Cs7g32410*, and *Cs2g20750* in sweet orange leaves infiltrated with a combination of Xcc*pthA4::*Tn5 strains carrying the pBBR5 constructs dTALECs2g27100, dTALECs7g18460, dTALEOrange1.1t00600, dTALECs9g17380, dTALCs6g17190, dTALECs7g32410, and dTALECs2g20750. Xcc*pthA4:*:Tn5 and Xcc*pthA4*::Tn5 dTALECsLOB1 served as negative and positive controls, respectively. Sweet orange leaves were infiltrated and collected at 7 dpi for gene expression assay using RT-qPCR. Error bars indicate means ± SEM (*n* = 3), and asterisks indicate significant differences in the expression of genes (∗*P* ≤ 0.05) compared to that of Xcc*pthA4:*:Tn5 mutant. The results shown are representative of three independent replicates.

**Fig. 4.**
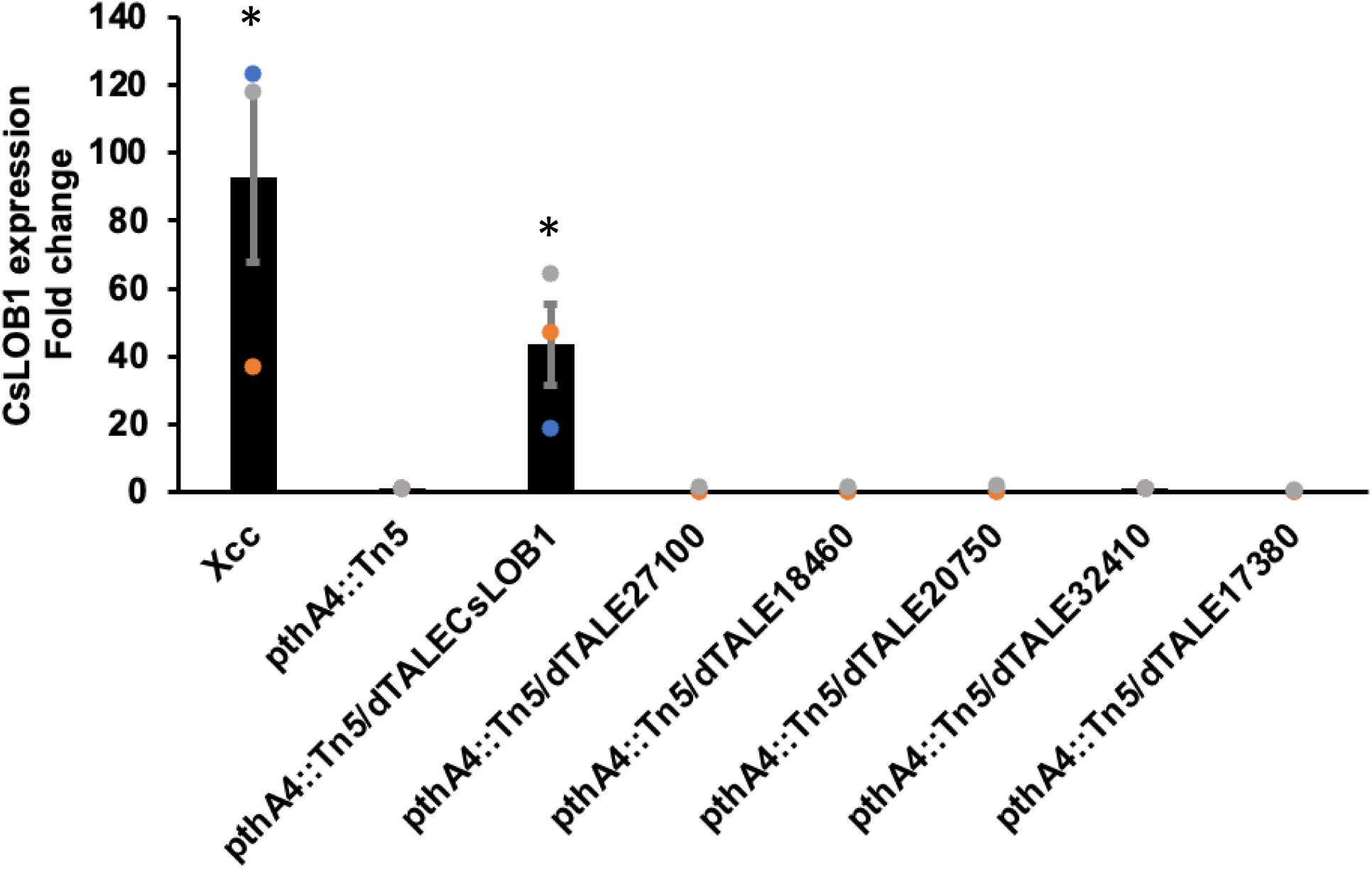
Expression analysis of *CsLOB1* in Sweet orange leaves infiltrated individually with Xcc or Xcc*PthA4::*Tn5 carrying the pBBR5 constructs for dTALECsLOB1, dTALECs2g27100, dTALECs7g18460, dTALECs2g20750, dTALECs7g32410, and dTALECs9g17380. *pthA4*::Tn5 served as negative control. Sweet orange leaves were infiltrated and collected at 2 dpi for gene expression assays using RT-qPCR. Error bars indicate means ± SEM (*n* = 3), and asterisks indicate significant differences in the expression of *CsLOB1* (∗*P* ≤ 0.05) compared to *pthA4:*:Tn5 mutant. The results shown are representative of three independent replicates.

To test whether dTALEs targeting the 7 CsLOB1 direct target genes can cause canker like symptoms, fully expanded young leaves of sweet orange were infiltrated with *Xcc* WT, *pthA4*::Tn5 mutant, dTALECsLOB1 and dTALEs at the concentration of OD600 of 0.2. First, we co-expressed a combination of 5 dTALEs (Cs2g20750, Cs6g17190, Cs7g32410, Cs7g18460, Orange1.1t00600) and 7 dTALEs (5 dTALEs plus dTALECs2g27100 and dTALECs9g17380) to comprehend if these dTALEs can cause canker like symptoms. Our results indicate that a combination of 5 and 7 dTALEs caused canker-like symptoms (Fig. 5). Next, we tested whether individual dTALEs can cause canker symptoms. Interestingly, dTALECs7g32410 and dTALECs2g20750 caused water soaking and pustules, dTALECs7g18460 and dTALECs6g17190 caused slightly yellowing, and the other three dTALEs did not cause any symptoms (Fig. 6).

**Fig. 5.**
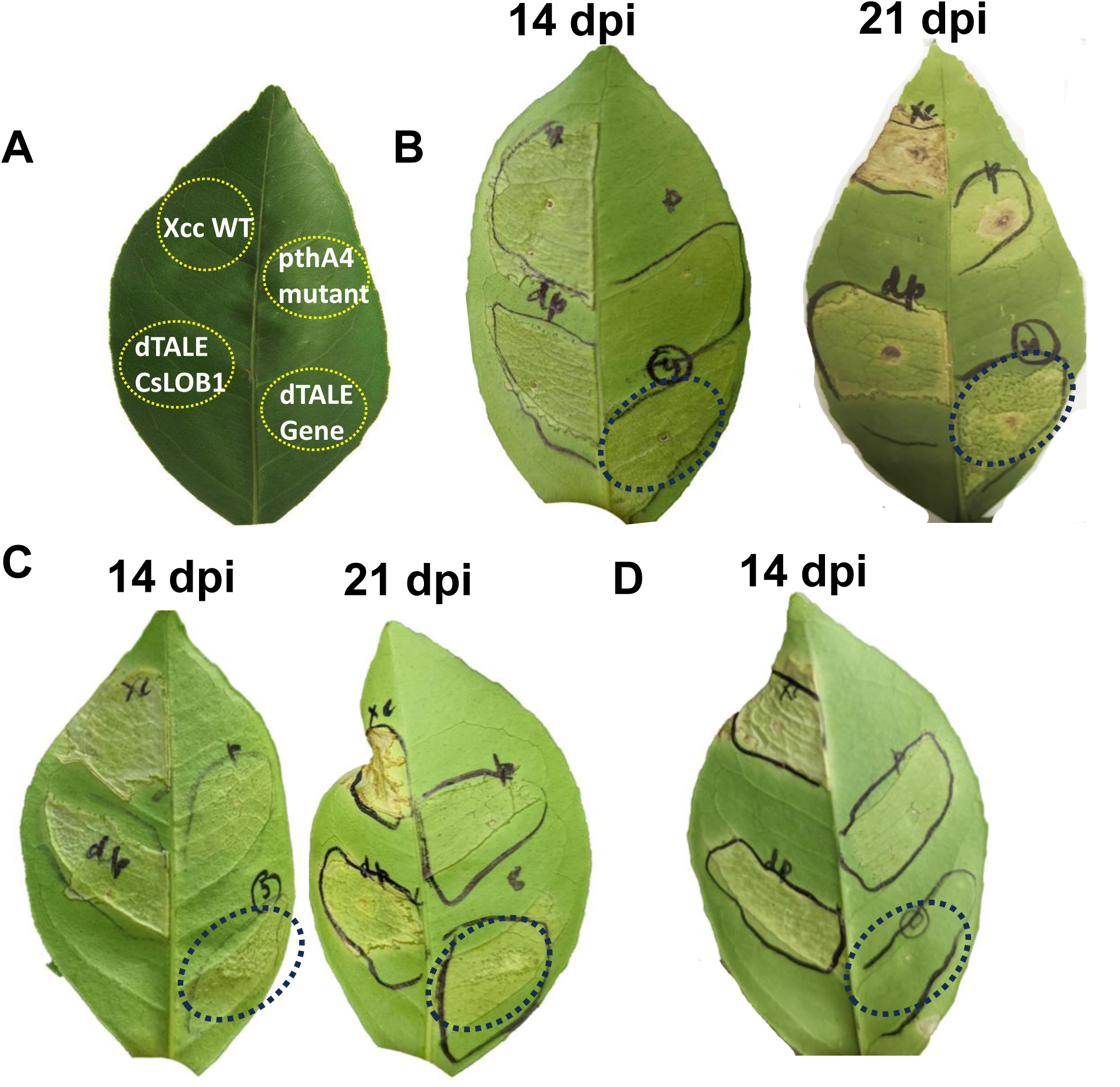
Virulence assay with constructed dTALEs in Sweet orange leaves: *Xcc* strains WT, Xcc*pthA4*::Tn5, Xcc*pthA4*::Tn5 dTALECsLOB1 and Xcc*pthA4*::Tn5 dTALEs for putative CsLOB1 target genes were infiltrated into fully expanded young leaves of Valencia sweet orange, which were evaluated for the symptoms at 14 and 21 dpi. Representative results were chosen from three independent experiments. A. Schematic representation of inoculation of leaves with different strains. B. Combination of 5 dTALEs shown with black dotted circle (Cs2g20750, Cs6g17190, Cs7g32410, Cs7g18460, Orange1.1t00600). C. Combination of 7 dTALEs shown with black dotted circle (Cs2g20750, Cs2g27100, Cs7g32410, Cs7g18460, Orange1.1t00600, Cs9g17380, Cs6g17190. D. mock control.

**Fig. 6.**
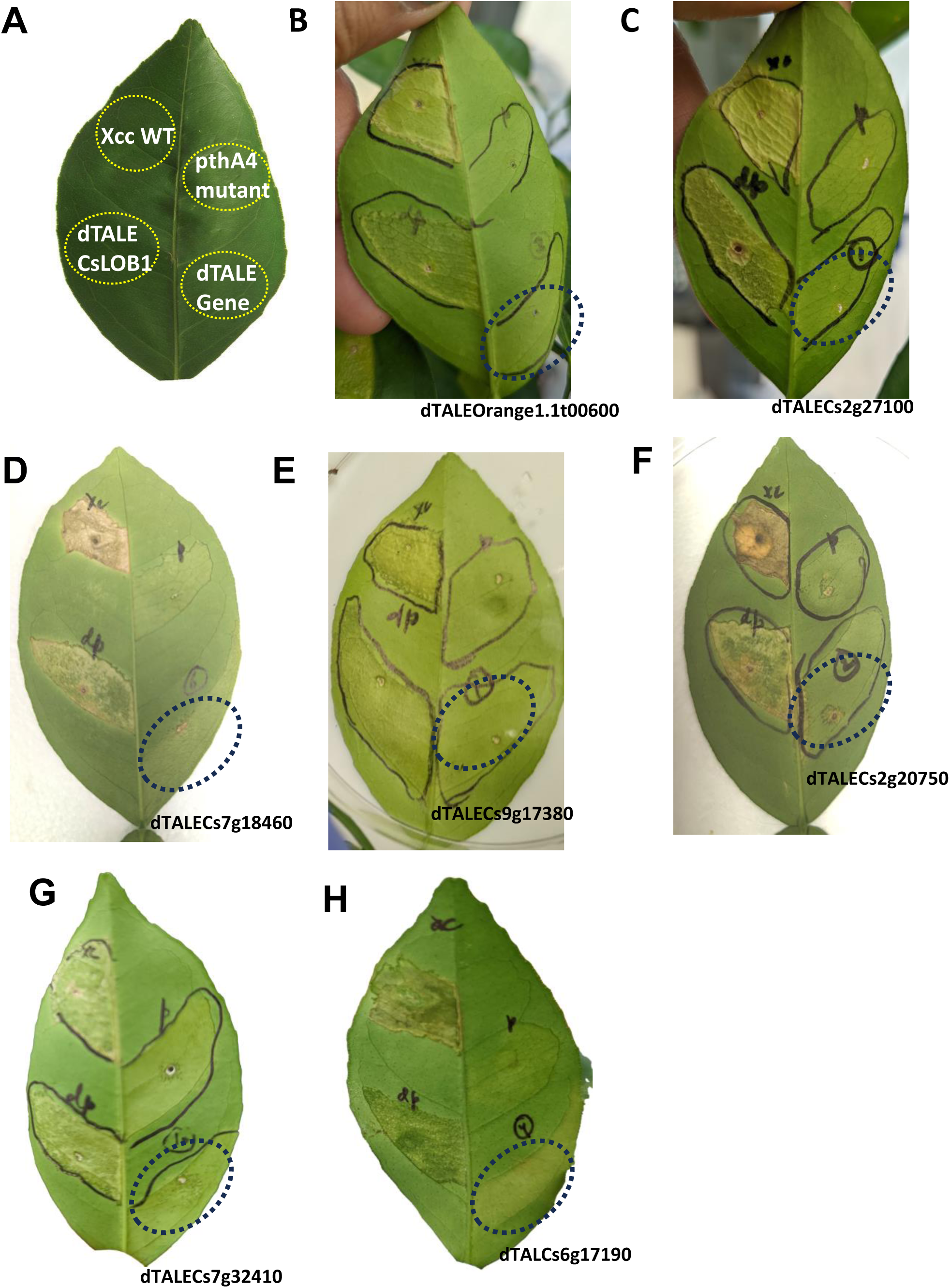
Virulence assay with dTALEs in Sweet orange leaves. *Xcc* strains WT, Xcc*pthA4*:: Tn5, Xcc*pthA4*::Tn5 dTALECsLOB1, and Xcc*pthA4*::Tn5 containing individual dTALE constructs were infiltrated into fully expanded Valencia sweet orange leaves (OD_600_ = 0.2). Representative results were chosen from three independent experiments. A. Schematic representation inoculation. B. dTALEOrange1.1t00600, C. dTALECs2g27100, D. dTALECs7g18460, E. dTALECs9g17380, F. dTALECs2g20750D, G. dTALECs7g32410, H. dTALCs6g17190.

## Discussion

In this study, we have demonstrated that either dTALECs7g32410 (CsEXP2) or dTALECs2g20750 (CsEG1) can cause water soaking and pustule symptoms, which are common phenomena during Xcc infection of citrus leaves [2]. dTALECs7g18460 and dTALECs6g17190 do not cause water soaking and pustules, but cause slightly yellowing. The canker like symptoms caused by either dTALECs7g32410 or dTALECs2g20750 individually are much less than the combined effect of 5 dTALEs (Cs2g20750, Cs6g17190, Cs7g32410, Cs7g18460, Orange1.1t00600) and 7 dTALEs (5 dTALEs plus dTALECs2g27100 and dTALECs9g17380), indicating a coordinated effect among CsEXP2 and CsEG1 or other 5 direct targets of CsLOB1 play incremental roles in canker symptom development. *CsEXP2* (Cs7g32410) encodes an expansin protein whereas *CsEG1* (Cs2g20750) encodes an endoglucanase protein. Expansins are non-enzymatic cell wall-loosening proteins. Expansins can disrupt non-covalent bonding between the cellulose microfibrils and cell wall matrix polysaccharides, which leads to extension of cell wall and cell expansion [39]. In contrast, endoglucanases enzymatically cleave β-1,4-glycosidic bonds such as those found in cellulose and xyloglucan [40]. Endoglucanases also play important roles in cell expansion. Xcc is known to cause enlarged cells (hypertrophy), a kind of cell expansion. Thus, we have provided direct evidence that *CsEXP2* (Cs7g32410) and *CsEG1* (Cs2g20750) are direct targets of CsLOB1 involved in canker symptom development. It is noteworthy that Cs2g20750 was identified to be a direct target of CsLOB1 via Chip-seq elsewhere [20]. Separation of cellulose microfibrils during cell expansion causes cell water uptake, thus triggering water soaking. Cs7g18460 encodes a glucomannan 4-beta-mannosyltransferase 11 whereas Cs6g17190 encodes a gibberellin-regulated protein 5. Glucomannan 4-beta-mannosyltransferases are involved in synthesis of galactomannan, a non-cellulosic polysaccharides of plant cell wall [41]. Gibberellin-regulated proteins have been reported to be involved in multiple plant growth and development processes [42]. How Cs7g18460 and Cs6g17190 contribute to canker symptom development remains to be explored.

Neither individual dTALEs nor mixed dTALEs can completely mimic the citrus canker symptoms caused by wild type Xcc in terms of the magnitude of water soaking, and pustule formation. Specifically, cell death was observed in the late stage of Xcc infection [43], but not in inoculation with dTALEs. This suggests other genes in addition to the 7 CsLOB1 direct targets are involved in citrus canker symptom development. For instance *CsLIEXP1* is a direct target of CsLOB1 and encodes another expansin gene [31]. It is probable that activation of *CsLIEXP1* contributes to hypertrophy phenomenon induced by Xcc. Multiple cell wall enzyme genes were reported to be direct targets of CsLOB1 including orange1.1t02719 (pectin esterase) and CsCEL20 (cellulase) [20, 33]. Pectate lyases, polygalacturonases, xyloglucan endotransglycosylase/hydrolase and other plant cell wall enzymes are involved in cell expansion [40]. Many genes encoding plant cell wall enzymes are positively regulated by CsLOB1 [20, 25, 34], thus contributing to hypertrophy and water soaking symptoms. Cs2g20600, which encodes a zinc finger C3HC4-type RING finger protein, is a direct target of CsLOB1 [30]. C3HC4-type RING finger proteins are E3 ubiquitin ligases, which might be involved in proteasomal degradation during cell death [44]. *CsNCED1-1* is a direct target of CsLOB1 and is a critical enzyme in ABA biosynthesis. ABA is a plant hormone that plays a crucial role in leaf senescence [45]. Activation of *CsNCED1-1* might contribute to the leaf drops caused by Xcc infection [43]. Interestingly dTALECsLOB1 causes very similar canker symptoms as wild type Xcc including water soaking, hypertrophy and hyperplasia except cell death, suggesting other factors induced by PthA4 might be responsible for the cell death development.

In sum, we have identified 7 direct targets of CsLOB1 and shown that Xcc indirectly activates *CsEXP2* and *CsEG1* via PthA4-CsLOB1 to cause canker symptoms. Identification of direct targets of CsLOB1 provides alternative target genes for disease resistance improvement via genome editing.

## Materials and methods

### Bacterial strains, plasmids, plant materials, and DNA manipulation

The bacterial strains and plasmids used in this study are listed in Supplemental Table 2. *E*. *coli* cells were cultured in Luria Bertani broth (LB) medium at 37°C. *Agrobacterium* strains were grown in LB containing rifampicin at 28°C. All *Xanthomonas citri* pv*. citri* (*Xcc)* mutant strains used in this study were derivatives of *Xcc* 306. *Xcc* strains were cultured in nutrient broth (NB) medium at 200 RPM or on nutrient agar (NA) plates at 28°C. When required, culture media were supplemented with appropriate antibiotics (kanamycin, 50 μg/mL; gentamicin, 10 μg/mL; spectinomycin, 100 μg/mL; rifampicin 50 μg/mL; and ampicillin, 100 μg/mL). *Nicotiana benthamiana* plants were cultivated in a growth chamber at 25°C with a 16 h light/8 h dark photoperiod. Four-to eight-week-old *N. benthamiana* plants were used for all the experiments. Citrus plants used in this study was Valencia sweet orange (*Citrus sinensis*). Plants were grown in a temperature-controlled (28°C) greenhouse under natural light conditions.

### Generation of GUS reporter and *CsLOB1* transient expression constructs

The binary vector pCAMBIA1380 was used to clone and express *CsLOB1* in *N. benthamiana*. pCAMBIA1380 contains the CaMV 35S promoter and a HA-tag downstream of the multiple cloning site, and the *CsLOB1* gene was cloned into the vector utilizing the *Xba*I and *Sal*I restriction sites. pCAMBIA construct was introduced into *Agrobacterium* strain EHA105 by freeze-thaw method [46]. The GUS reporter system was used to assess the transcriptional activation of select target genes by LOB1. Promoter sequences located ∼1 kb upstream of the translational start site in Cs1g16560, Cs7g32410, Cs6g17190, Cs2g27100, Cs9g17380, Cs2g27090, Cs2g21440, Cs7g18460, Cs5g33680, orange1.1t00600, Cs2g05560, Cs2g20750, and Cs2g20600 were cloned into the binary GUS reporter construct pCAMBIA1380 35S-GUS using primers listed in Supplemental Table 3. Promoter constructs were transformed into *Agrobacterium* strain EHA105. *Agrobacterium* transformants were cultured in LB medium overnight. Cultures were harvested by centrifugation (4000 g for 10 min), washed, and resuspended in infiltration buffer (10 mM MgCl2, 0.2 mM acetosyringone and 200 mM MES, pH 5.6) to a final OD_600_=0.5. Buffer-supplemented *Agrobacterium* strains were incubated at room temperature for 2-4 h, then the *CsLOB1* and reporter constructs were co-infiltrated (OD_600_ = 1.0) into leaves of 4-7 weeks old *N. benthamiana* with needleless syringes for transient expression assays [31]. In quantitative assays, three leaf discs (1 cm) were collected at 2 dpi, and GUS activity was measured using 4-methylumbelliferyl-β-glucuronide described previously [47]. Proteins were quantified using the Bradford method. The experiment was repeated three times with three biological replicates for each treatment.

### Construction of dTALEs

dTALEs were assembled using “Golden Gate TALEN and TAL Effector Kit 2.0” as previously described [48] and cloned into pTAL2 as a final destination vector. A library of four basic repeats encoding RVDs NG, NI, HD, and NH, which correspond to target nucleotides T, A, C, and G, respectively, were used. The repeat regions of artificial dTALEs were assembled using the RVDs corresponding to the targeted nucleotides near the TATA box in the promoter regions of Cs7g32410, Cs6g17190, Cs2g27100, Cs9g17380, Cs7g18460, orange1.1t00600, Cs2g20750, Orange1.1t00817, Cs5g34710, and Cs5g06600 (Fig. 2A). The broad host destination vector pBBRNPth with the −23 to +444 N-terminal coding fragment of *pthA4* and an HA tag cloned into pBBR1MCS-5 [43] was used for subcloning the dTALEs RVDs along with N- and C-terminal regions. The pTAL2 *Pst*I/*Eco*RI fragments containing the dTALEs were cloned into pBBRNPth for expression in *Xanthomonas*, and dTALE sequences were validated by PCR, digestion, and sequencing. Detailed information on the dTALEs, including RVDs and targeted EBE sequences, is shown in Fig.2A. All constructs were introduced into *Xcc pthA4*::Tn5 by electroporation.

### Detection and confirmation of dTALE through western blot analysis

The expression of dTALEs was confirmed by western blot using mouse anti-HA antibody. *Xcc pthA4* mutant strains containing dTALEs constructs were killed in sodium-dodecyl sulfate-polyacrylamide gel electrophoresis (SDS-PAGE) loading buffer (100 mM Tris-HCl, pH 6.8; 9% β-mercapto ethanol, 40% glycerol, 0.0005% bromophenol blue, 4% SDS) via heating for 10 min at 95°C, which were subjected to gel electrophoresis afterwards. Separated proteins were transferred onto the nitrocellulose membrane (Biorad, USA). Proteins were detected by an anti-HA antibody (Sigma Aldrich, USA), and developed on the X-ray film using kit from (Thermo Scientific, USA)

### dTALEs delivery into plant cells via *Xanthomonas*

dTALEs were delivered into plant cells via *XccpthA4*::Tn5. The pBBRNpth-dTALEs plasmids (Supplementary Table 2), containing an HA-tag epitope in the c-terminus of dTALEs, were electroporated (2.5 kv, 5 ms) into *X*cc*pthA4*::Tn5 strain. The *Xcc* strains - WT, *pthA4*::Tn5 mutant and mutants containing dTALEs for targeting the gene of interest and, mutant containing dTALE for targeting *CsLOB1* were cultured overnight in NA liquid medium with respective antibiotics. The bacterial cells were collected by centrifugation (5000 g, 10 min), washed twice with 10 mM MgCl_2_ and, resuspended in 10 mM MgCl_2_ to OD_600_ = 0.2. The suspensions were infiltrated into fully expanded young leaves of Valencia sweet orange with needleless syringes. Infiltration of simply 10 mM buffer was used as a mock. The leaf phenotype was photographed at 14-21 dpi. These experiments were repeated three times with similar results.

### Plant inoculations, and quantification of bacterial populations

Bacterial suspensions (OD_600_=0.2) for monitoring symptom development, RNA isolation and physiological measurements, were syringe-infiltrated into fully expanded young leaves of two- to four-year-old Valencia sweet orange plants. After inoculations, plants were kept in a temperature-controlled (28°C) greenhouse under natural light conditions.

### RNA isolation and reverse transcription-quantitative PCR (RT-qPCR)

RNA was isolated from leaf tissues inoculated with *Xcc* WT, *pthA4*::Tn5 mutant, *pthA4*::Tn5 (dTALEpthA4), and *pthA4*::Tn5 carrying dTALEs targeting different genes. For each sample, three 1-cm-diameter leaf disks from inoculated areas were pooled from three leaves belonging to the same plant and frozen in liquid nitrogen immediately. Samples were ground into fine powder using TissueLyser II (QIAGEN, Hilden, Germany). Total RNA was extracted using RNeasy plant RNA isolation kit (Thermo Fisher Scientific, Waltham, MA, USA). For RT-qPCR, 1 µg RNA samples were treated with Qiagen DNA out solution and reverse-transcribed using a qScript cDNA Synthesis Kit (Qiagen, Germany). cDNAs were amplified with gene-specific primer (Supplementary Table 3) using SYBR Green master mix (Thermo Fisher Scientific, USA) by the Quant Studio 3 Real-Time PCR System (Applied Biosystems Inc., Foster City, CA). The *GAPDH* gene was used as an endogenous control for normalization, and gene expression level was calculated by the comparative *Ct* method [49].

## Acknowledgments

We thank Wang lab members for constructive suggestions and insightful discussions. This project was supported by funding from Florida Citrus Initiative Program, FDACS Specialty Crop Block Grant Program AM22SCBPFL1125, and Hatch project [FLA-CRC-005979] (N. Wang).

---

**Supplementary Table 1.** Genes positively regulated by PthA4 and CsLOB1.

| Genes | CsLOB1 overexpression vs wild type | WT vs *pthA4* mutant grapefruit | Annotation |
| --- | --- | --- | --- |
| Cs1g16560 | 1.86 | 1.545646 | Pectin esterase |
| Cs7g32410 | 4.46 | 2.868992 | Expansin A4 |
| Cs6g17190 | 8.08 | 1.860488 | Gibberellin-regulated protein 5 |
| Cs2g27100 | 3.99 | 1.907112 | GDSL esterase/lipase |
| Cs9g17380 | 5 | 3.631467 | Putative uncharacterized protein |
| Cs2g27090 | 6.47 | 0.969382 | GDSL esterase/lipase At5g14450 |
| Cs2g21440 | 3.84 | 2.718401 | Putative uncharacterized protein |
| Cs3g18550 | 1.68 | 1.847577 | Pollen-specific protein C13 |
| Cs7g18460 | 5.28 | 2.562127 | glucomannan 4-beta-mannosyltransferase 2 |
| Cs5g33680 | 2.55 | 1.154933 | Glycosyltransferase |
| orange1.1t00600 | 1.96 | 1.691821 | 3-oxo-5-alpha-steroid 4-dehydrogenase |
| Cs2g05560 | 6.48 | 2.342324 | Protein E6 |
| Cs2g20750 | 4.15 | 4.752483 | Endoglucanase 11 |

Note: Numbers depict Log<sub>2</sub> fold change.

**Supplementary Table 2.** Strains or plasmids used in this study.

| Strains | Relevant characteristics | Source |
| --- | --- | --- |
| *Xanthomonas citri* subsp. *citri* |  |  |
| Xcc306 | wild type | [1] |
| Xcc306 *pthA4*::Tn5 | *pthA4* deletion mutant with Tn5 insertion, Kan<sup>r</sup> | [1] |
| *pthA4*::Tn5/dTALE<sub>pthA4</sub> | Artificial TALE targeting CsLOB1 complement Xcc306 *pthA4*::Tn5, Kan<sup>r</sup> Gm<sup>r</sup> | [2] |
| *pthA4*::Tn5/dTALECs9g17380 | Artificial TALE targeting Cs9g17380, Gm<sup>r</sup> | This study |
| *pthA4*::Tn5/dTALECs2g27100 | Artificial TALE targeting Cs2g27100, Kan<sup>r</sup> Gm<sup>r</sup> | This study |
| *pthA4*::Tn5/dTALEOrange1.1t00600 | Artificial TALE targeting Orange1.1t00600, Kan<sup>r</sup> Gm<sup>r</sup> | This study |
| *pthA4::Tn5/dTALECs2g20750* | Artificial TALE targeting Cs2g20750, Kan<sup>r</sup> Gm<sup>r</sup> | This study |
| *pthA4::Tn5/dTALECs7g32410* | Artificial TALE targeting Cs7g32410, Kan<sup>r</sup> Gm<sup>r</sup> | This study |
| *pthA4::Tn5/dTALECs7g18460* | Artificial TALE targeting Cs7g18460, Kan<sup>r</sup> Gm<sup>r</sup> | This study |
| *pthA4::Tn5/dTALECs6g17190* | Artificial TALE targeting Cs6g17190, Kan<sup>r</sup> Gm<sup>r</sup> | This study |
| *pthA4::Tn5/dTALECs5g06600* | Artificial TALE targeting Cs5g06600, Kan<sup>r</sup> Gm<sup>r</sup> | This study |
| *pthA4::Tn5/dTALECs5g34710* | Artificial TALE targeting Cs5g34710, Kan<sup>r</sup> Gm<sup>r</sup> | This study |
| *pthA4::Tn5/dTALEOrange1.1t00817* | Artificial TALE targeting Orange1.1t00817, Kan<sup>r</sup> Gm<sup>r</sup> | This study |
| *E. coli* |  |  |
| Stellar | HST04 strain, dam-/dcn- | Invitrogen |
| *Agrobacterium tumefaciens* |  |  |
| EHA105 | C58, pTiBo542DT-DNA, Rif<sup>r</sup> | [3] |
| Plasmids |  |  |
| erGFP-pCAMBIA 1380N-35S:HA | Binary vector, Kan<sup>r</sup> | www.cambia.org |
| pCAMBIA1380-35SGUS1 | Binary vector containing promoterless gusA; used for reporter assays, Kan<sup>r</sup> | www.cambia.org |
| pBBR1MCS-5 | Broad host expression vector. Gm<sup>R</sup> | [4] |
| pBBRNPth | pBBR1MCS-5 derivative for expression of -24 to +444 N-terminal coding fragment of PthA4 (XACb0065) and a HA tag. Used as backbone for construction of dTALEs. Gm<sup>R</sup> | [2] |
| pTAL2 | Intermediate destination vector for assembled dTALEs. Amp<sup>R</sup> | [5] |
| dTALECs9g17380 | pBBR1MCS-5 expressing designed TALE targeting Cs9g17380 fused to HA tag. Gm<sup>R</sup> | This study |
| dTALECs2g27100 | pBBR1MCS-5 expressing designed TALE targeting Cs7g27100 fused to HA tag. Gm<sup>R</sup> | This study |
| dTALEOrange1.1t00600 | pBBR1MCS-5 expressing designed TALE targeting Orange1.1t00600 fused to HA tag. Gm<sup>R</sup> | This study |
| dTALECs2g20750 | pBBR1MCS-5 expressing designed TALE targeting Cs2g20750 fused to HA tag. Gm<sup>R</sup> | This study |
| dTALECs7g32410 | pBBR1MCS-5 expressing designed TALE targeting Cs7g32410 fused to HA tag. Gm<sup>R</sup> | This study |
| dTALECs6g17190 | pBBR1MCS-5 expressing designed TALE targeting Cs6g17190 fused to HA tag. Gm<sup>R</sup> | This study |
| dTALECs5g06600 | pBBR1MCS-5 expressing designed TALE targeting Cs5g06600 fused to HA tag. Gm<sup>R</sup> | This study |
| dTALECs5g34710 | pBBR1MCS-5 expressing designed TALE targeting Cs5g34710 fused to HA tag. Gm<sup>R</sup> | This study |
| dTALEOrange1.1t00817 | pBBR1MCS-5 expressing designed TALE targeting Orange1.1t00817 fused to HA tag. Gm<sup>R</sup> | This study |
| 35S:CsLOB1-HA | erGFP-pCAMBIA 1380N expressing CsLOB1 under 35S promoter and fused to HA tag | This study |
| pCs7g32410::uidA | ~1 kb promoter region of Cs7g32410 cloned upstream of uidA for GUS expression in pCAMBIA1380 | This study |
| pOrange1.1t00600:: uidA | ~1 kb promoter region of Orange1.1t00600 cloned upstream of uidA for GUS expression in pCAMBIA1380 | This study |
| pCs5g33680:: uidA | ~1 kb promoter region of Cs5g33680 cloned upstream of uidA for GUS expression in pCAMBIA1380 | This study |
| pCs2g21440:: uidA | ~1 kb promoter region of Cs2g21440 cloned upstream of uidA for GUS expression in pCAMBIA1380 | This study |
| pCs2g27090::*uidA* | ~1 kb promoter region of Cs2g27090 cloned upstream of *uidA* for GUS expression in pCAMBIA1380 | This study |
| pCs7g18460::*uidA* | ~1 kb promoter region of Cs7g18460 cloned upstream of *uidA* for GUS expression in pCAMBIA1380 | This study |
| pCs2g20600::*uidA* | ~1 kb promoter region of Cs2g20600 cloned upstream of *uidA* for GUS expression in pCAMBIA1380 | This study |
| pCs9g17380::*uidA* | ~1 kb promoter region of Cs9g17380 cloned upstream of *uidA* for GUS expression in pCAMBIA1380 | This study |
| pCs2g27100::*uidA* | ~1 kb promoter region of Cs2g27100 cloned upstream of *uidA* for GUS expression in pCAMBIA1380 | This study |
| pCs6g17190::*uidA* | ~1 kb promoter region of Cs6g17190 cloned upstream of *uidA* for GUS expression in pCAMBIA1380 | This study |
| pCs5g33680::*uidA* | ~1 kb promoter region of Cs5g33680 cloned upstream of *uidA* for GUS expression in pCAMBIA1380 | This study |
| pOrange1.1t00600::*uidA* | ~1 kb promoter region of Orange1.1t00600 cloned upstream of *uidA* for GUS expression in pCAMBIA1380 | This study |

**Supplementary Table 3.**
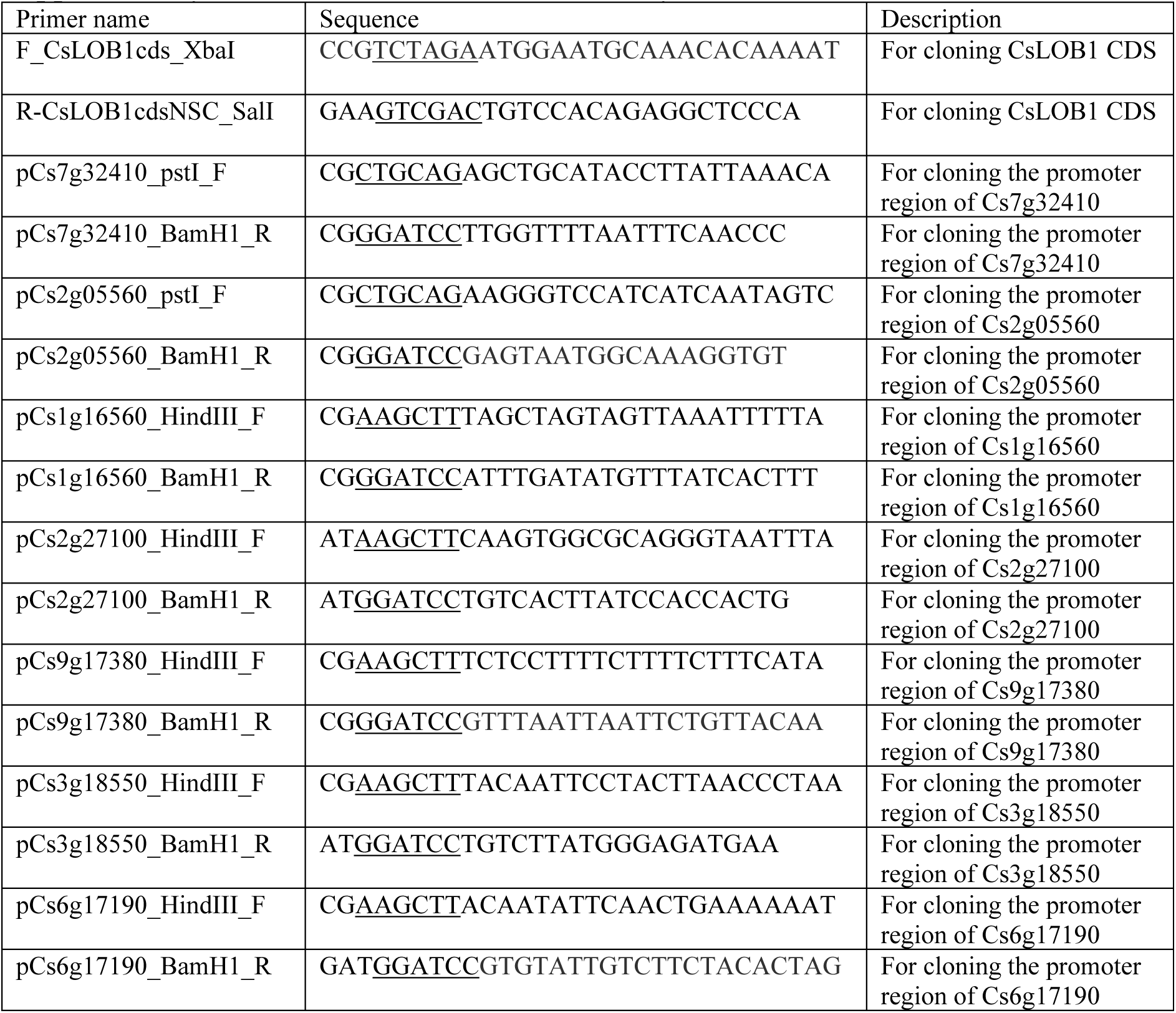
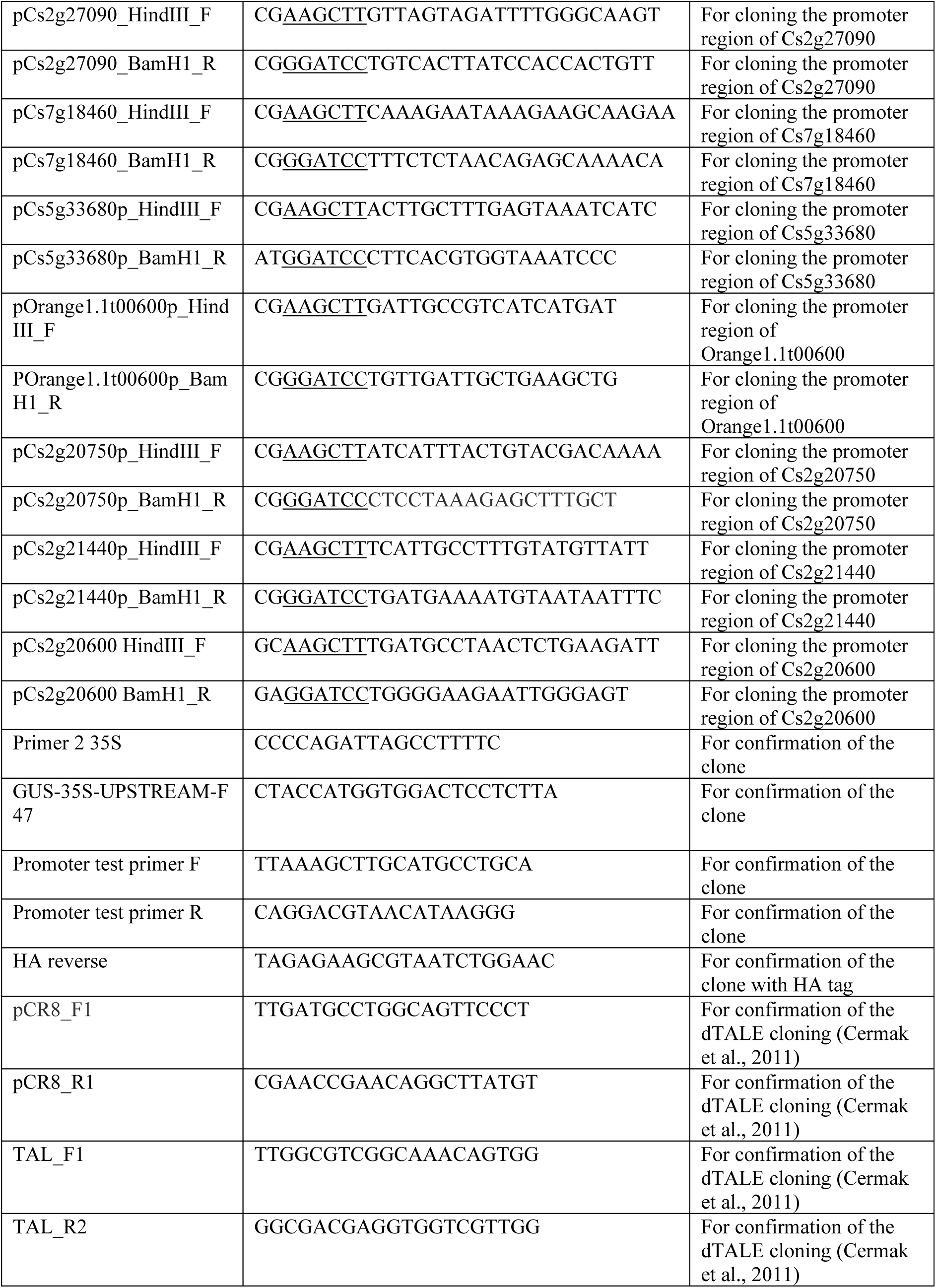
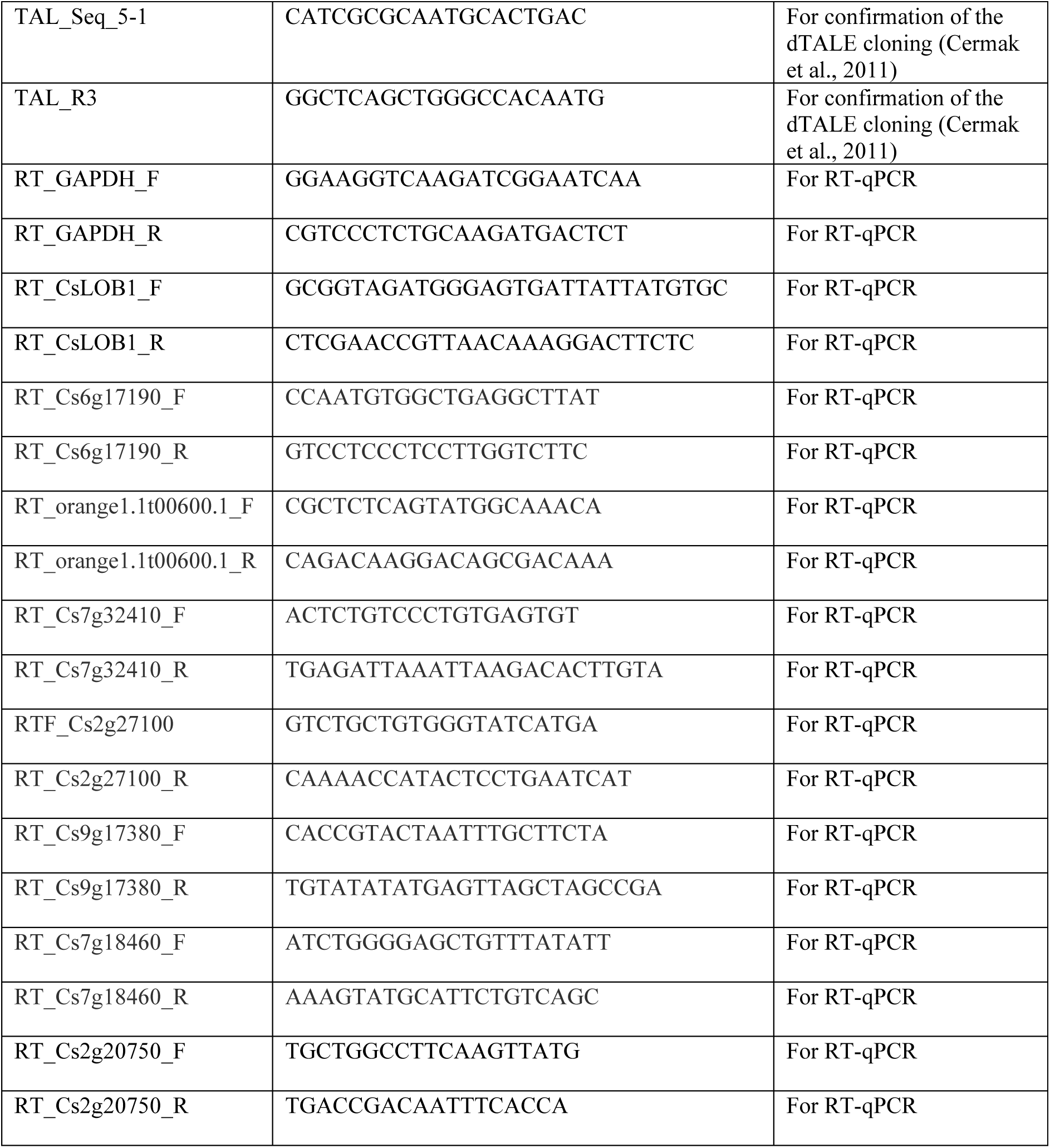
Primers used in this study.

## Literature cited

1. Ference CM, Gochez AM, Behlau F, Wang N, Graham JH, Jones JB. Recent advances in the understanding of Xanthomonas citri ssp. citri pathogenesis and citrus canker disease management. Mol Plant Pathol. 2018;19(6):1302–18. Epub 2018/03/08. doi: 10.1111/mpp.12638. PubMed PMID: 29105297.

2. Brunings AM, Gabriel DW. Xanthomonas citri: breaking the surface. Mol Plant Pathol. 2003;4(3):141–57. doi: 10.1046/j.1364-3703.2003.00163.x. PubMed PMID: 20569374.

3. Yan Q, Wang N. High-throughput screening and analysis of genes of Xanthomonas citri subsp. citri involved in citrus canker symptom development. Mol Plant Microbe Interact. 2011. doi: 10.1094/MPMI-05-11-0121. PubMed PMID: 21899385.

4. Hu Y, Zhang J, Jia H, Sosso D, Li T, Frommer WB, et al. Lateral organ boundaries 1 is a disease susceptibility gene for citrus bacterial canker disease. Proc Natl Acad Sci U S A. 2014;111(4):E521-9. doi: 10.1073/pnas.1313271111. PubMed PMID: 24474801; PubMed Central PMCID: PMCPMC3910620.

5. Tang X, Wang X, Huang Y, Ma L, Jiang X, Rao MJ, et al. Natural variations of TFIIAγ gene and LOB1 promoter contribute to citrus canker disease resistance in Atalantia buxifolia. PLoS Genet. 2021;17(1):e1009316. Epub 2021/01/26. doi: 10.1371/journal.pgen.1009316. PubMed PMID: 33493197; PubMed Central PMCID: PMCPMC7861543.

6. Cernadas RA, Doyle EL, Niño-Liu DO, Wilkins KE, Bancroft T, Wang L, et al. Code-assisted discovery of TAL effector targets in bacterial leaf streak of rice reveals contrast with bacterial blight and a novel susceptibility gene. PLoS Pathog. 2014;10(2):e1003972. doi: 10.1371/journal.ppat.1003972. PubMed PMID: 24586171; PubMed Central PMCID: PMCPMC3937315.

7. Kay S, Hahn S, Marois E, Hause G, Bonas U. A bacterial effector acts as a plant transcription factor and induces a cell size regulator. Science. 2007;318(5850):648-51. doi: 10.1126/science.1144956. PubMed PMID: 17962565.

8. Zhou J, Peng Z, Long J, Sosso D, Liu B, Eom JS, et al. Gene targeting by the TAL effector PthXo2 reveals cryptic resistance gene for bacterial blight of rice. Plant J. 2015. doi: 10.1111/tpj.12838. PubMed PMID: 25824104.

9. Cohn M, Bart RS, Shybut M, Dahlbeck D, Gomez M, Morbitzer R, et al. Xanthomonas axonopodis virulence is promoted by a transcription activator-like effector-mediated induction of a SWEET sugar transporter in cassava. Mol Plant Microbe Interact. 2014;27(11):1186–98. doi: 10.1094/mpmi-06-14-0161-r. PubMed PMID: 25083909.

10. Oliva R, Ji C, Atienza-Grande G, Huguet-Tapia JC, Perez-Quintero A, Li T, et al. Broad-spectrum resistance to bacterial blight in rice using genome editing. Nat Biotechnol. 2019;37(11):1344–50. Epub 2019/10/28. doi: 10.1038/s41587-019-0267-z. PubMed PMID: 31659337; PubMed Central PMCID: PMCPMC6831514.

11. Li T, Liu B, Spalding MH, Weeks DP, Yang B. High-efficiency TALEN-based gene editing produces disease-resistant rice. Nat Biotechnol. 2012;30(5):390–2. doi: 10.1038/nbt.2199. PubMed PMID: 22565958.

12. Huang X, Wang Y, Wang N. Highly Efficient Generation of Canker-Resistant Sweet Orange Enabled by an Improved CRISPR/Cas9 System. Front Plant Sci. 2021;12:769907. Epub 20220111. doi: 10.3389/fpls.2021.769907. PubMed PMID: 35087548; PubMed Central PMCID: PMCPMC8787272.

13. Jia H, Zhang Y, Orbović V, Xu J, White FF, Jones JB, et al. Genome editing of the disease susceptibility gene CsLOB1 in citrus confers resistance to citrus canker. Plant Biotechnol J. 2017;15(7):817–23. Epub 2017/01/04. doi: 10.1111/pbi.12677. PubMed PMID: 27936512; PubMed Central PMCID: PMCPMC5466436.

14. Peng A, Chen S, Lei T, Xu L, He Y, Wu L, et al. Engineering canker-resistant plants through CRISPR/Cas9-targeted editing of the susceptibility gene CsLOB1 promoter in citrus. Plant Biotechnol J. 2017;15(12):1509–19. Epub 2017/04/04. doi: 10.1111/pbi.12733. PubMed PMID: 28371200; PubMed Central PMCID: PMCPMC5698050.

15. Jia H, Orbovic V, Jones JB, Wang N. Modification of the PthA4 effector binding elements in Type I CsLOB1 promoter using Cas9/sgRNA to produce transgenic Duncan grapefruit alleviating XccΔpthA4:dCsLOB1.3 infection. Plant Biotechnol J. 2016;14(5):1291–301. doi: 10.1111/pbi.12495. PubMed PMID: 27071672.

16. Jia H, Omar AA, Orbović V, Wang N. Biallelic Editing of the LOB1 Promoter via CRISPR/Cas9 Creates Canker-Resistant ‘Duncan’ Grapefruit. Phytopathology®. 2021;112(2):308–14. doi: 10.1094/PHYTO-04-21-0144-R.

17. Jia H, Wang Y, Su H, Huang X, Wang N. LbCas12a-D156R Efficiently Edits LOB1 Effector Binding Elements to Generate Canker-Resistant Citrus Plants. Cells. 2022;11(3). doi: 10.3390/cells11030315.

18. Jia H, Omar AA, Xu J, Dalmendray J, Wang Y, Feng Y, et al. Generation of transgene-free canker-resistant Citrus sinensis cv. Hamlin in the T0 generation through Cas12a/CBE co-editing. Front Plant Sci. 2024;15:1385768. Epub 20240326. doi: 10.3389/fpls.2024.1385768. PubMed PMID: 38595767; PubMed Central PMCID: PMCPMC11002166.

19. Huang X, Wang Y, Wang N. Base Editors for Citrus Gene Editing. Front Genome Ed. 2022;4:852867. Epub 20220228. doi: 10.3389/fgeed.2022.852867. PubMed PMID: 35296063; PubMed Central PMCID: PMCPMC8919994.

20. Zou X, Du M, Liu Y, Wu L, Xu L, Long Q, et al. CsLOB1 regulates susceptibility to citrus canker through promoting cell proliferation in citrus. Plant J. 2021;106(4):1039–57. Epub 2021/03/24. doi: 10.1111/tpj.15217. PubMed PMID: 33754403.

21. de Lima LFF, Carvalho IGB, de Souza-Neto RR, Dos Santos LDS, Nascimento CA, Takita MA, et al. Antisense Oligonucleotide as a New Technology Application for CsLOB1 Gene Silencing Aiming at Citrus Canker Resistance. Phytopathology. 2024;114(8):1802–9. Epub 20240814. doi: 10.1094/phyto-02-24-0058-kc. PubMed PMID: 38748545.

22. Huang X, Jia H, Xu J, Wang Y, Wen J, Wang N. Transgene-free genome editing of vegetatively propagated and perennial plant species in the T0 generation via a co-editing strategy. Nat Plants. 2023;9(10):1591–7. Epub 20230918. doi: 10.1038/s41477-023-01520-y. PubMed PMID: 37723203.

23. Su H, Wang Y, Xu J, Omar AA, Grosser JW, Calovic M, et al. Generation of the transgene-free canker-resistant Citrus sinensis using Cas12a/crRNA ribonucleoprotein in the T0 generation. Nat Commun. 2023;14(1):3957. Epub 20230705. doi: 10.1038/s41467-023-39714-9. PubMed PMID: 37402755; PubMed Central PMCID: PMCPMC10319737.

24. Su H, Wang Y, Xu J, Omar AA, Grosser J, Wang N. Cas12a/3 crRNAs RNP transformation enables transgene-free multiplex genome editing, long deletions, and inversions in citrus chromosome in the T0 generation. bioRxiv. 2024:2024.06.13.598908. doi: 10.1101/2024.06.13.598908.

25. Zhang J, Huguet-Tapia JC, Hu Y, Jones J, Wang N, Liu S, et al. Homologues of CsLOB1 in citrus function as disease susceptibility genes in citrus canker. Mol Plant Pathol. 2017;18(6):798–810. Epub 2016/08/11. doi: 10.1111/mpp.12441. PubMed PMID: 27276658.

26. Xu C, Luo F, Hochholdinger F. LOB Domain Proteins: Beyond Lateral Organ Boundaries. Trends Plant Sci. 2016;21(2):159–67. doi: 10.1016/j.tplants.2015.10.010. PubMed PMID: 26616195.

27. Majer C, Hochholdinger F. Defining the boundaries: structure and function of LOB domain proteins. Trends Plant Sci. 2011;16(1):47–52. doi: 10.1016/j.tplants.2010.09.009. PubMed PMID: 20961800.

28. Fan M, Xu C, Xu K, Hu Y. LATERAL ORGAN BOUNDARIES DOMAIN transcription factors direct callus formation in Arabidopsis regeneration. Cell Res. 2012;22(7):1169–80. Epub 20120417. doi: 10.1038/cr.2012.63. PubMed PMID: 22508267; PubMed Central PMCID: PMCPMC3391013.

29. Kim MJ, Kim M, Lee MR, Park SK, Kim J. LATERAL ORGAN BOUNDARIES DOMAIN (LBD)10 interacts with SIDECAR POLLEN/LBD27 to control pollen development in Arabidopsis. Plant J. 2015;81(5):794–809. doi: 10.1111/tpj.12767. PubMed PMID: 25611322.

30. Duan S, Jia H, Pang Z, Teper D, White F, Jones J, et al. Functional characterization of the citrus canker susceptibility gene CsLOB1. Molecular plant pathology. 2018;19(8):1908–16. doi: 10.1111/mpp.12667. PubMed PMID: 29461671.

31. de Souza-Neto RR, Vasconcelos FNdC, Teper D, Carvalho Isis Gabriela B, Takita MA, Benedetti CE, et al. The Expansin Gene CsLIEX P1 Is a Direct Target of CsLOB1 in Citrus. Phytopathology®. 2023;113(7):1266–77. doi: 10.1094/PHYTO-11-22-0424-R.

32. Chen X, Zou H, Zhuo T, Rou W, Wu W, Fan X. Xanthomonas citri subsp. citri type III effector PthA4 directs the dynamical expression of a putative citrus carbohydrate-binding protein gene for canker formation. Elife. 2024;13. Epub 20240813. doi: 10.7554/eLife.91684. PubMed PMID: 39136681; PubMed Central PMCID: PMCPMC11321762.

33. Li Y, Lou H, Fu H, Su H, Hao C, Luo J, et al. Identifying the role of cellulase gene CsCEL20 upon the infection of Xanthomonas citri subsp. citri in citrus. Mol Breed. 2025;45(1):10. Epub 20250106. doi: 10.1007/s11032-024-01531-3. PubMed PMID: 39781329; PubMed Central PMCID: PMCPMC11704107.

34. Hu Y, Duan S, Zhang Y, Shantharaj D, Jones JB, Wang N. Temporal Transcription Profiling of Sweet Orange in Response to PthA4-Mediated Xanthomonas citri subsp. citri Infection. Phytopathology. 2016;106(5):442–51. doi: 10.1094/PHYTO-09-15-0201-R. PubMed PMID: 26780431.

35. Bell EM, Lin WC, Husbands AY, Yu L, Jaganatha V, Jablonska B, et al. Arabidopsis lateral organ boundaries negatively regulates brassinosteroid accumulation to limit growth in organ boundaries. Proc Natl Acad Sci U S A. 2012;109(51):21146–51. doi: 10.1073/pnas.1210789109. PubMed PMID: 23213252; PubMed Central PMCID: PMCPMC3529045.

36. Husbands A, Bell EM, Shuai B, Smith HM, Springer PS. LATERAL ORGAN BOUNDARIES defines a new family of DNA-binding transcription factors and can interact with specific bHLH proteins. Nucleic Acids Res. 2007;35(19):6663–71. doi: 10.1093/nar/gkm775. PubMed PMID: 17913740; PubMed Central PMCID: PMCPMC2095788.

37. Schwartz AR, Morbitzer R, Lahaye T, Staskawicz BJ. TALE-induced bHLH transcription factors that activate a pectate lyase contribute to water soaking in bacterial spot of tomato. Proc Natl Acad Sci U S A. 2017;114(5):E897-E903. Epub 2017/01/18. doi: 10.1073/pnas.1620407114. PubMed PMID: 28100489; PubMed Central PMCID: PMCPMC5293091.

38. Morbitzer R, Elsaesser J, Hausner J, Lahaye T. Assembly of custom TALE-type DNA binding domains by modular cloning. Nucleic Acids Res. 2011;39(13):5790–9. doi: 10.1093/nar/gkr151. PubMed PMID: 21421566; PubMed Central PMCID: PMCPMC3141260.

39. Cosgrove DJ. Assembly and enlargement of the primary cell wall in plants. Annu Rev Cell Dev Biol. 1997;13:171–201. doi: 10.1146/annurev.cellbio.13.1.171. PubMed PMID: 9442872.

40. Glass M, Barkwill S, Unda F, Mansfield SD. Endo-β-1,4-glucanases impact plant cell wall development by influencing cellulose crystallization. J Integr Plant Biol. 2015;57(4):396–410. doi: 10.1111/jipb.12353. PubMed PMID: 25756224.

41. Liepman AH, Wilkerson CG, Keegstra K. Expression of cellulose synthase-like (Csl) genes in insect cells reveals that CslA family members encode mannan synthases. Proc Natl Acad Sci U S A. 2005;102(6):2221–6. Epub 20050112. doi: 10.1073/pnas.0409179102. PubMed PMID: 15647349; PubMed Central PMCID: PMCPMC548565.

42. Inomata N. Gibberellin-regulated protein allergy: Clinical features and cross-reactivity. Allergology International. 2020;69(1):11–8. doi: 10.1016/j.alit.2019.10.007.

43. Teper D, Xu J, Li J, Wang N. The immunity of Meiwa kumquat against Xanthomonas citri is associated with a known susceptibility gene induced by a transcription activator-like effector. PLoS Pathog. 2020;16(9):e1008886. Epub 2020/09/16. doi: 10.1371/journal.ppat.1008886. PubMed PMID: 32931525; PubMed Central PMCID: PMCPMC7518600.

44. Bader M, Steller H. Regulation of cell death by the ubiquitin-proteasome system. Curr Opin Cell Biol. 2009;21(6):878–84. Epub 20091021. doi: 10.1016/j.ceb.2009.09.005. PubMed PMID: 19850458; PubMed Central PMCID: PMCPMC2818673.

45. Jakubowicz M, Nowak W, Gałgański Ł, Babula-Skowrońska D, Kubiak P. Expression profiling of the genes encoding ABA route components and the ACC oxidase isozymes in the senescing leaves of Populus tremula. J Plant Physiol. 2020;248:153143. Epub 20200226. doi: 10.1016/j.jplph.2020.153143. PubMed PMID: 32126452.

46. Walkerpeach CR, Velten J. Agrobacterium-mediated gene transfer to plant cells: cointegrate and binary vector systems. In: Gelvin SB, Schilperoort RA, editors. Plant Molecular Biology Manual. Dordrecht: Springer Netherlands; 1994. p. 33-51.

47. Jia H, Wang N. Xcc-facilitated agroinfiltration of citrus leaves: a tool for rapid functional analysis of transgenes in citrus leaves. Plant Cell Rep. 2014;33(12):1993–2001. doi: 10.1007/s00299-014-1673-9. PubMed PMID: 25146436.

48. Teper D, Wang N. Consequences of adaptation of TAL effectors on host susceptibility to Xanthomonas. PLoS Genet. 2021;17(1):e1009310. Epub 2021/01/20. doi: 10.1371/journal.pgen.1009310. PubMed PMID: 33465093; PubMed Central PMCID: PMCPMC7845958.

49. Pfaffl MW. A new mathematical model for relative quantification in real-time RT-PCR. Nucleic Acids Res. 2001;29(9):e45. Epub 2001/05/09. doi: 10.1093/nar/29.9.e45. PubMed PMID: 11328886; PubMed Central PMCID: PMCPMC55695.

## References

[1] Yan Q, Wang N. High-throughput screening and analysis of genes of Xanthomonas citri subsp. citri involved in citrus canker symptom development. Mol Plant Microbe Interact. 2011. doi: 10.1094/MPMI-05-11-0121. PubMed PMID: 21899385.

[2] Teper D, Wang N. Consequences of adaptation of TAL effectors on host susceptibility to Xanthomonas. PLoS Genet. 2021;17(1):e1009310. Epub 2021/01/20. doi: 10.1371/journal.pgen.1009310. PubMed PMID: 33465093; PubMed Central PMCID: PMCPMC7845958.

[3] Wirawan IG, Kang HW, Kojima M. Isolation and characterization of a new chromosomal virulence gene of Agrobacterium tumefaciens. J Bacteriol. 1993;175(10):3208–12. doi: 10.1128/jb.175.10.3208-3212.1993. PubMed PMID: 8491736; PubMed Central PMCID: PMCPMC204646.

[4] Kovach ME, Phillips RW, Elzer PH, Roop RM, 2nd, Peterson KM. pBB R1M CS: a broad-host-range cloning vector. Biotechniques. 1994;16(5):800–2. PubMed PMID: 8068328.

[5] Cermak T, Doyle EL, Christian M, Wang L, Zhang Y, Schmidt C, et al. Efficient design and assembly of custom TALEN and other TAL effector-based constructs for DNA targeting. Nucleic Acids Res. 2011;39(12):e82. doi: 10.1093/nar/gkr218. PubMed PMID: 21493687; PubMed Central PMCID: PMCPMC3130291.

